# Prenatal psychosocial stress-induced behavioral and neuroendocrine abnormalities are associated with sex-specific alterations in synaptic transmission and differentially modulated by maternal environment

**DOI:** 10.1101/2020.05.20.106674

**Authors:** Sandra P. Zoubovsky, Michael T. Williams, Sarah Hoseus, Shivani Tumukuntala, Amy Riesenberg, Jay Schulkin, Charles V. Vorhees, Kenneth Campbell, Hee-Woong Lim, Louis J. Muglia

**Author notes:** Corresponding author: Louis J. Muglia MD PhD, Cincinnati Children’s Hospital Medical Center, 240 Albert Sabin Way, Room S11.334, Cincinnati, OH 45229.

## Abstract

Prenatal stress (PS) is associated with increased vulnerability to affective disorders. Transplacental glucocorticoid passage and stress-induced maternal environment alterations are recognized as potential routes of transmission that can fundamentally alter neurodevelopment. However, molecular mechanisms underlying aberrant emotional outcomes or the individual contributions intrauterine stress versus maternal environment play in shaping these mechanisms remain unknown. Here, we report anxiogenic behaviors, anhedonia, and female hypothalamic-pituitary-adrenal axis hyperactivity as a consequence of psychosocial PS in mice. Sex-specific placental responses to stress and evidence of fetal amygdala programming precede these abnormalities. In adult offspring, we observe amygdalar transcriptional changes demonstrating sex-specific dysfunction in synaptic transmission and neurotransmitter systems. We find these abnormalities are primarily driven by in-utero stress exposure. Importantly, maternal care changes postnatally reverse anxiety-related behaviors and partially rescue gene alterations associated with neurotransmission. Our data demonstrate the influence maternal environment exerts in shaping offspring emotional development despite deleterious effects of intrauterine stress.

## INTRODUCTION

Substantial evidence from human and animal studies indicates that exposure to prenatal stress (PS) is a critical risk factor for developing neuropsychiatric disorders later in life (Bale, 2015; Bale et al., 2010; Faa et al., 2016). However, the contributing mechanisms by which these intrauterine challenges program disease susceptibility remain largely unknown. Studies in rodents have demonstrated that PS can result in the emergence of anxiety-like and depressive-related behaviors, reduced social interaction, and deficits in attention and learning (Brunton, 2013; Weinstock, 2015). These phenotypes are often accompanied by dysregulation in hypothalamic-pituitary-adrenal (HPA) axis activity (Glover et al., 2010), the main neuroendocrine system regulating responses to stress, and such dysregulation is a common feature in humans suffering from depression and anxiety disorders (Young, 2004). The effects of PS in rodents on offspring neurodevelopmental outcomes seem to be in part dependent on the type of stress experienced, timing of exposure, and offspring sex (Bale, 2015; Brunton, 2013; Leshem & Schulkin, 2012). PS paradigms commonly utilized in animal studies rely on physical stressors, such as restraint, or do not accurately portray the multifaceted nature of stress experienced by women (Bleker et al., 2019; Weinstock, 2016). As such, it becomes imperative to study the effects of gestational insults that are more translationally relevant such as exposure to chronic variable psychosocial stressors.

Human and rodent studies have also demonstrated that PS is correlated with abnormalities in maternal behavior (Hillerer et al., 2012). There is substantial evidence indicating that variations in maternal care can strongly influence offspring behavior and HPA axis activity (Bale, 2015; Hackman et al., 2010). This raises the intriguing question of whether the neurodevelopmental programming effects of maternal stress are a consequence of in-utero disruptions or alterations in maternal care.

The programming effects of PS on offspring behavior and neuroendocrine function seem to be in part mediated by changes in circulating maternal glucocorticoids, the primary stress hormone secreted by the HPA axis (Bale, 2015; Bale et al., 2010; Seckl & Holmes, 2007). During pregnancy, the placenta plays a critical role in maintaining in-utero homeostasis and tightly regulates maternal-fetal glucocorticoid transfer through the actions of placental glucocorticoid barrier proteins (Bronson & Bale, 2016; Jansson & Powell, 2007; Seckl & Holmes, 2007). The 11β-hydroxysteroid dehydrogenase type 2 (11ß–HSD2) and type 1 (11ß-HSD1) enzymes degrade most circulating glucocorticoids into inactive metabolites and catalyze the reverse reaction, respectively (Chapman et al., 2013). P-glycoprotein (ABCB1) serves as a membrane-bound efflux protein that prevents the transfer of xenobiotic substances and glucocorticoids to the fetus (Mark et al., 2009). Increasing evidence from animal studies suggests that placental function can be affected by maternal adversity resulting in changes in 11ß– HSD2 expression in a temporal and offspring sex-specific manner. For example, PS during early pregnancy results in decreased 11ß– HSD2 levels in female offspring and a trend toward increased expression in male offspring, but mid to late gestational stress is associated with a reduction in both sexes (Jensen Peña et al., 2012; Lesage et al., 2001; Pankevich et al., 2009). Despite these advances, the sex-specific alterations in the various components of the placental glucocorticoid barrier in response to chronic psychosocial insults and its influence on neurodevelopment remain unknown.

PS alters the developmental trajectory of vulnerable brain structures, resulting in functional changes that are thought to underlie the risk for developing emotional disorders (Bale, 2015; Harris & Seckl, 2011). The amygdala is a key site for integrating neuroendocrine and behavioral responses to stress and therefore plays an essential role in emotion regulation (Arnett et al., 2015). To date, most research examining the effects of PS paradigms that are psychosocial in nature (exposure to social defeat) have focused on measuring gene expression changes in key molecular regulators of the HPA axis (Brunton, 2013). Up-regulation in the glucocorticoid receptor (GR) and corticotropin releasing hormone (CRH) has been noted, as well as altered CRH receptor (CRH R) levels (Brunton & Russell, 2010; Mueller & Bale, 2008; Zohar & Weinstock, 2011). While these findings underscore the importance of characterizing changes in gene transcription in the amygdala to better understand the physiological basis of affective disturbances, sex-specific alterations in amygdalar transcriptional profiles in response to psychosocial PS and whether these changes arise in-utero or from alterations in maternal care remains to be investigated.

To address these questions, we developed a chronic gestational stress (CGS) paradigm which consists of exposing pregnant mice to psychosocially challenging insults presented in an unpredictable fashion from gestational day 6.5 – 17.5. Employing these mild to moderate psychosocial manipulations allows us to better mimic stressors experienced by women during pregnancy (Yim et al., 2015). We have previously found exposure to our paradigm results in the development of depression and anxiety-like phenotypes in dams as well as abnormalities in maternal behavior, evidenced by the emergence of fragmented and erratic maternal care patterns (Zoubovsky et al., 2020). Here, we investigated the effects of psychosocial PS on placental glucocorticoid barrier components as well as offspring behavior and neuroendocrine function. To elucidate the molecular mechanisms underlying the various phenotypes observed, we compared amygdalar transcriptional profiles of control (CTRL) and PS offspring since the amygdala represents a nodal point between the HPA and emotional output. Lastly, to dissociate the impact of in-utero stress from abnormalities in maternal care resulting from stress during gestation, we performed cross-fostering (CF) of pups at birth between CTRL and CGS dams (see Figure 1 for schematic outlining of the experimental paradigm).

**Figure 1.**
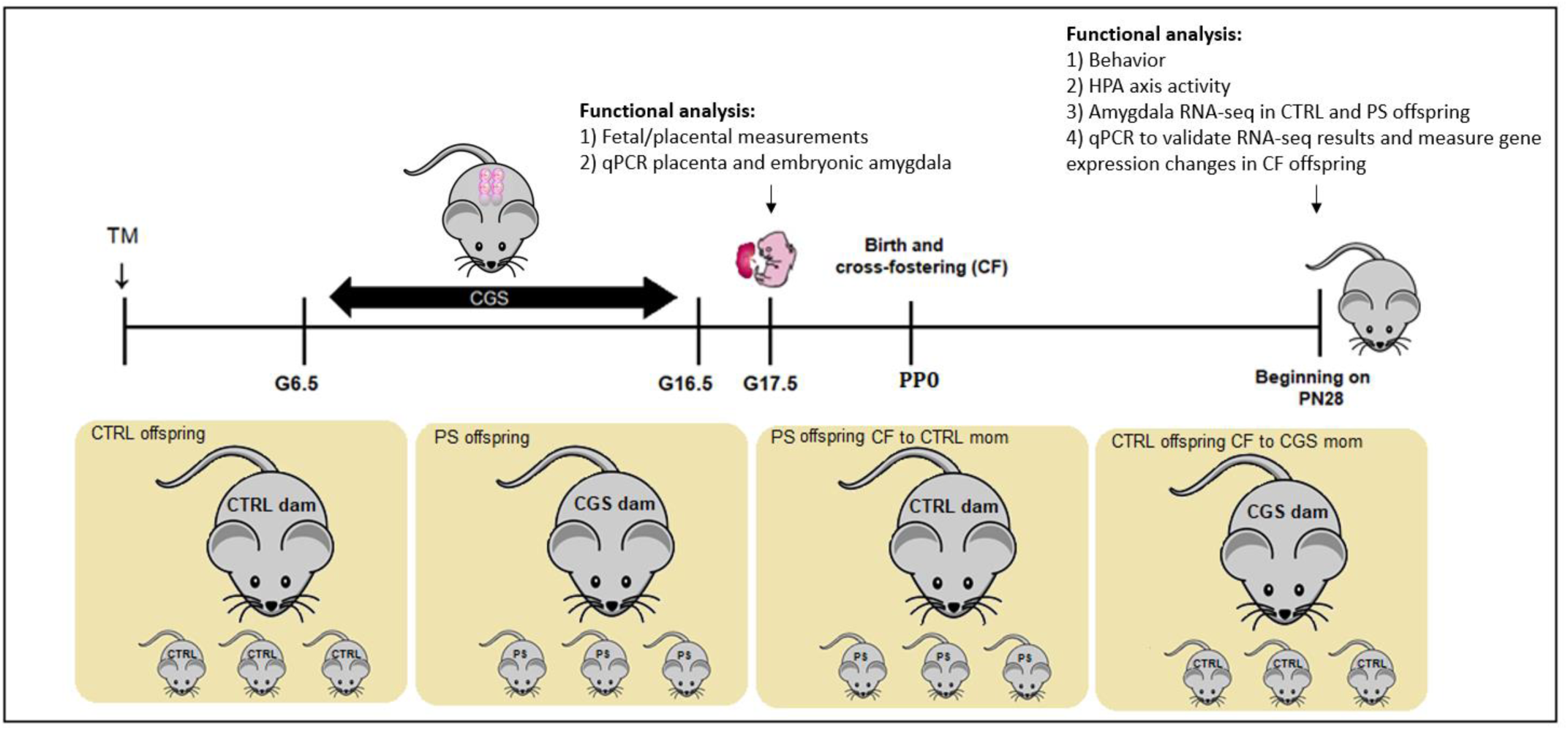
Experimental design. Psychosocial stress was performed from gestational day 6.5 (G6.5) to G17.5. Fetal and placental measurements and qPCR on placental glucocorticoid barrier components was performed on G17.5. A separate subset of mice was cross-fostered (CF) at birth so as to generate 4 separate groups for our studies. Functional analysis was carried out in these four groups beginning at postnatal (PN) day 28, including behavior and neuroendocrine characterization. RNA-seq analysis was performed in control and PS offspring amygdala samples and a subset of differentially expressed genes from this analysis were measured in cross-fostered offspring amygdala samples via qPCR.

## RESULTS

### Psychosocial PS leads to anxiety-related behaviors, an increased state of alertness, and anhedonia, phenotypes that are differentially affected by the postnatal maternal environment

We first measured the effects of psychosocial PS on anxiety-like behaviors via light dark transition box (LD). A significant PS x CF interaction was detected in the total amount of time spent in the light zone of LD (Figure 2A; F_1,27.7_= 9.68, p=0.0043). Offspring exposed to psychosocial PS showed a decrease in total time spent in the light zone when compared with age-matched CTRL offspring (P=0.0023). This reduction was reversed in PS offspring CF to CTRL moms (P=0.0005), suggesting the emergence of an anxiety-like phenotype following PS that is significantly improved by changes in maternal care. No differences were observed between CTRL offspring and CTRL offspring CF to CGS moms (P=0.6456), indicating the fragmented maternal care patterns displayed by dams exposed to psychosocial stress during pregnancy (Zoubovsky et al., 2020) is not contributing to the emergence of anxiety-related behaviors in offspring. When evaluating the total number of entries into the light zone, a significant CF effect was detected (Figure 2B; F_1,19.1_= 40.44, P<0.0001), where both PS offspring raised by CTRL mothers and CTRL offspring raised by CGS dams had more entries into the light zone. Despite these changes, analysis of overall locomotion in an open field test (OFT) revealed no significant effect of CF (Figure 2C; F_1,26.9_= 0.04, P=0.8419) or PS effect (Figure 2C; F_1,26.9_= 0.48, P=0.4940), suggesting that behavioral abnormalities or normalization of anxiety-like behaviors in PS offspring CF to CTRL mothers are not due to ambulatory changes.

**Figure 2.**
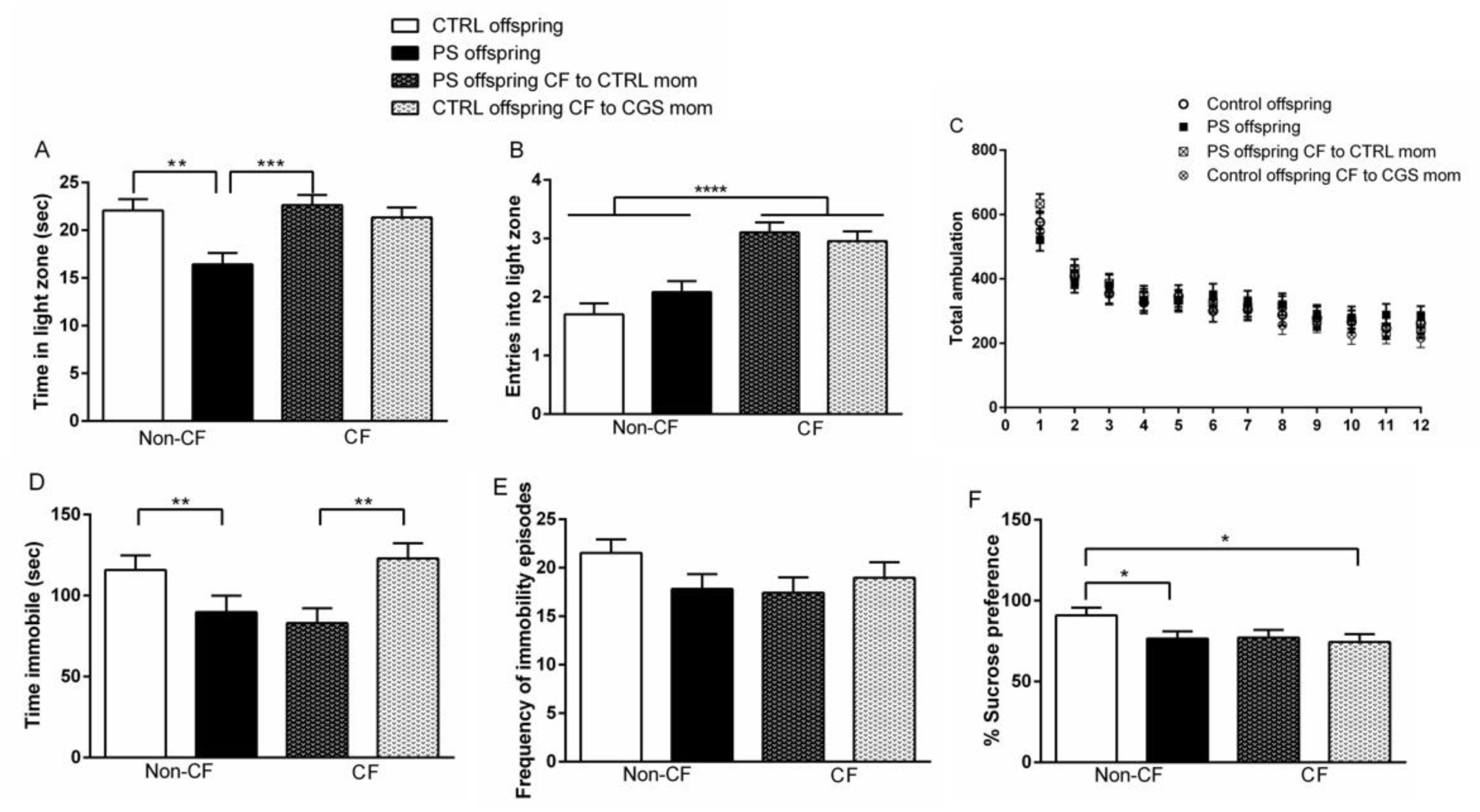
Psychosocial PS leads to behavioral abnormalities in both male and female offspring that are differentially modulated by postnatal maternal environment. (**A**) Time spent and (**B**) entries into light compartment in LD, CTRL offspring = 30, PS offspring = 30, PS offspring CF to CTRL mom = 39, CTRL offspring CF to CGS mom = 40. (**C**) Total ambulation in OFT, CTRL offspring = 30, PS offspring = 30, PS offspring CF to CTRL mom = 39, CTRL offspring CF to CGS mom = 40. (**D**) Time spent immobile and (**E**) frequency of immobility episodes in FST, CTRL offspring = 38, PS offspring = 31, PS offspring CF to CTRL mom = 30, CTRL offspring CF to CGS mom = 29. (**F**) Percent sucrose preference as measured in the SPT, CTRL offspring = 28, PS offspring = 30, PS offspring CF to CTRL mom = 28, CTRL offspring CF to CGS mom = 28. Data presented as mean ± SEM. \**p* < 0.05, \*\**p* < 0.01, \*\*\**p* < 0.001, \*\*\*\**p* < 0.0001 mixed linear ANOVA with prenatal stress x cross-fostering x sex model and litter as a randomized block factor. For OFT, interval was a repeated measures factor.

In a separate experiment, we assessed responsiveness to stress in the forced swim test (FST). We found a significant PS effect on the total amount of time spent immobile (Figure 2D; F_1,30.6_= 12.12, P=0.0015). Interestingly, both PS offspring and PS offspring CF to CTRL mothers exhibited a significant reduction in immobility time when compared with respective CTRLs. We interpret this measure to perhaps reflect an increased state of alertness when faced with a stressor, in this case swimming, which could be associated with a higher vulnerability for aberrant emotional outcomes, such as anxiety and/or impulsivity. Our data further suggest this heightened alertness is a result of in-utero stress and not modulated by changes in postnatal environment (e.g. maternal care) as time spent immobile in CTRL offspring raised by CGS dams was similar to CTRL offspring raised by CTRL mothers (Figure 2D; F_1,30.6_= 0.54, P=0.4673 for PS x CF interaction). No differences were observed across groups in the total number of immobility episodes (Figure 2E; F_1,25_= 2.91, P=0.1006 for PS effect; F_1,25_= 0.91, P=0.3487 for CF effect). We used the sucrose preference test (SPT) to evaluate anhedonia, a symptom often seen in depression (Planchez et al., 2019). A significant PS x CF interaction was observed on the preference for 4% sucrose solution (Figure 2F; F_1,26.9_= 3.24, p=0.0416). When allowed to freely choose between water or 4% sucrose solution, PS offspring displayed significantly reduced preference for 4% sucrose than age-matched CTRL offspring (P=0.0396), a measure considered to represent an inability to experience pleasure. Interestingly, the emergence of this anhedonic behavior seemed to be mediated by both effects of in-utero stress and alterations in postnatal maternal environment as CTRL offspring raised by stressed mothers also exhibited a reduction in sucrose preference when compared with non-CF CTRLs (P=0.0221). Our data suggest that anhedonic characteristics could not be rescued by normalizing maternal care delivered to pups as PS offspring CF to CTRL dams exhibited a similar reduction in sucrose preference as non-CF PS offspring did (P=0.9252). Noteworthy, no significant sex x PS interaction was found on the amount of time spent in the light zone in LD (Figure 2A; F_1,105_= 0.00, P=0.9951), on the amount of time spent immobile in the FST (Figure 2D; F_1,93.9_= 1.62, P=0.2064), and on 4% sucrose preference in the SPT (Figure 2F; F_1, 84.9_= 0.00, P=0.9816), indicating male and female PS offspring were equally affected in the behavioral parameters measured by these assays.

There were no differences in the total amount of time spent socializing with a stranger mouse in the social interaction assay (SI) in PS offspring when compared with CTRLs (Figure 2 – figure supplement 1A) or in associative learning as measured by no changes in freezing behavior during the training phase or context and cued testing phases of fear conditioning (FC) (Figure 2 – figure supplement 1B-D).

### Psychosocial PS leads to acute stress-induced HPA axis hyperactivity in female offspring, an effect not influenced by maternal environmental changes

In order to determine the effects of psychosocial PS on neuroendocrine function, we measured circadian concentrations in serum corticosterone (CORT) levels and in response to a novel acute stressor at postnatal day (PN) 28. No differences were observed in CORT measurements at the circadian nadir timepoint (Figure 3A; F_1,21_= 2.92, P=0.1023 for PS effect; F_1,21_= 0.03, P=0.8598 for sex effect; F_1,21_= 0.22, P=0.6462 for PS x sex interaction). Analysis of peak CORT values revealed no effect of PS (Figure 3B; F_1,20.3_= 0.00, P=0.9631), although there was a significant main effect of sex (Figure 3B; F_1,22_= 198.13, p<0.0001). Increased CORT was measured at the peak timepoint in females across all our groups when compared with males, a normal sex-variation often noted in HPA axis activity (Rao & Androulakis, 2017). Following 15-min of swimming, there was a significant PS x sex interaction on serum CORT levels (Figure 3C; F_1,105_= 6.39, P=0.0129). Elevation in stress-induced CORT secretion was observed in PS females when compared with PS males (P=0.0012). Importantly, PS females and PS females CF to CTRL dams had similar serum CORT levels in response to the 15-min swim, which were significantly higher than CTRL females and CF CTRL females (P<0.0001). These data suggest that in females, psychosocial PS results in HPA axis hyperresponsivity following acute stress exposure and this effect is primarily modulated by in-utero stress as elevated CORT release was also observed in PS females CF to CTRL dams.

**Figure 3.**
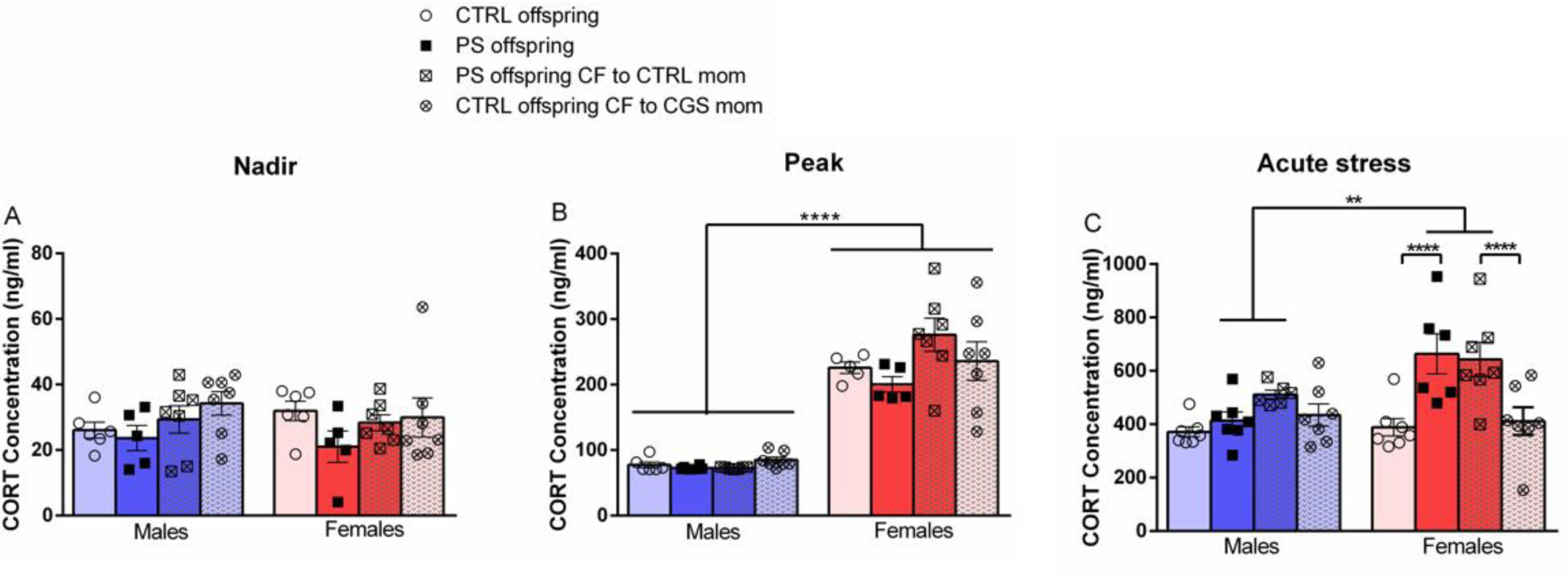
Psychosocial PS leads to HPA axis dysregulation in female offspring that are not reversed by changes in postnatal maternal care. Serum CORT measurements performed on PN28 at (**A**) nadir, (**B**) peak, and (**C**) after 15 min swim, CTRL offspring = 5-7, PS offspring = 5-7, PS offspring CF to CTRL mom = 6-7, CTRL offspring CF to CGS mom = 7, per sex per time point. Data presented as mean ± SEM. \*\**p* < 0.01, \*\*\*\**p* < 0.0001 mixed linear ANOVA with prenatal stress x cross-fostering x sex model and litter as a randomized block factor.

### Psychosocial PS induces sex-specific placental responses and is associated with developmental programming of the fetal amygdala

To evaluate the effects of psychosocial PS on placental function, we measured fetal and placental weights and calculated the fetal to placental weight ratio at gestational day 17.5 in CTRL and PS mice. We observed a significant main effect of sex on fetal weight (Figure 4A; F_1,64.2_= 13.54, P=0.0005), where female fetuses were lighter than male fetuses. However, no significant PS effect was noted (F_1,7_= 0.06, P=0.8163). For placental weights, there was a significant main effect of sex (Figure 4B; F_1,69.1_= 12.88, P=0.0006) and a significant interaction of sex x PS (Figure 4B; F_1,69.1_= 9.20, P=0.003). Placentas from female fetuses that were exposed to psychosocial PS were significantly lighter than placentas from CTRL females (P=0.0036) or PS males (P<0.0001). No differences were found between CTRL male and female placentas (P=0.6872). Together, these data suggest placentas from females have a potential to compensate for environmental insults as they are able to maintain weight of female fetuses after PS exposure despite the reduction observed in placental weights. Consistent with this notion, significant sex effect (Figure 4C; F_1,65_= 4.78, P=0.0325) and sex x PS interaction (Figure 4C; F_1,65_= 6.56, P=0.0128) were observed on the fetal to placental weight ratio. This ratio was significantly elevated in PS females when compared with PS males (P=0.0017) and there was a trend for it to be increased in PS females than CTRL females (P=0.0996).

**Figure 4.**
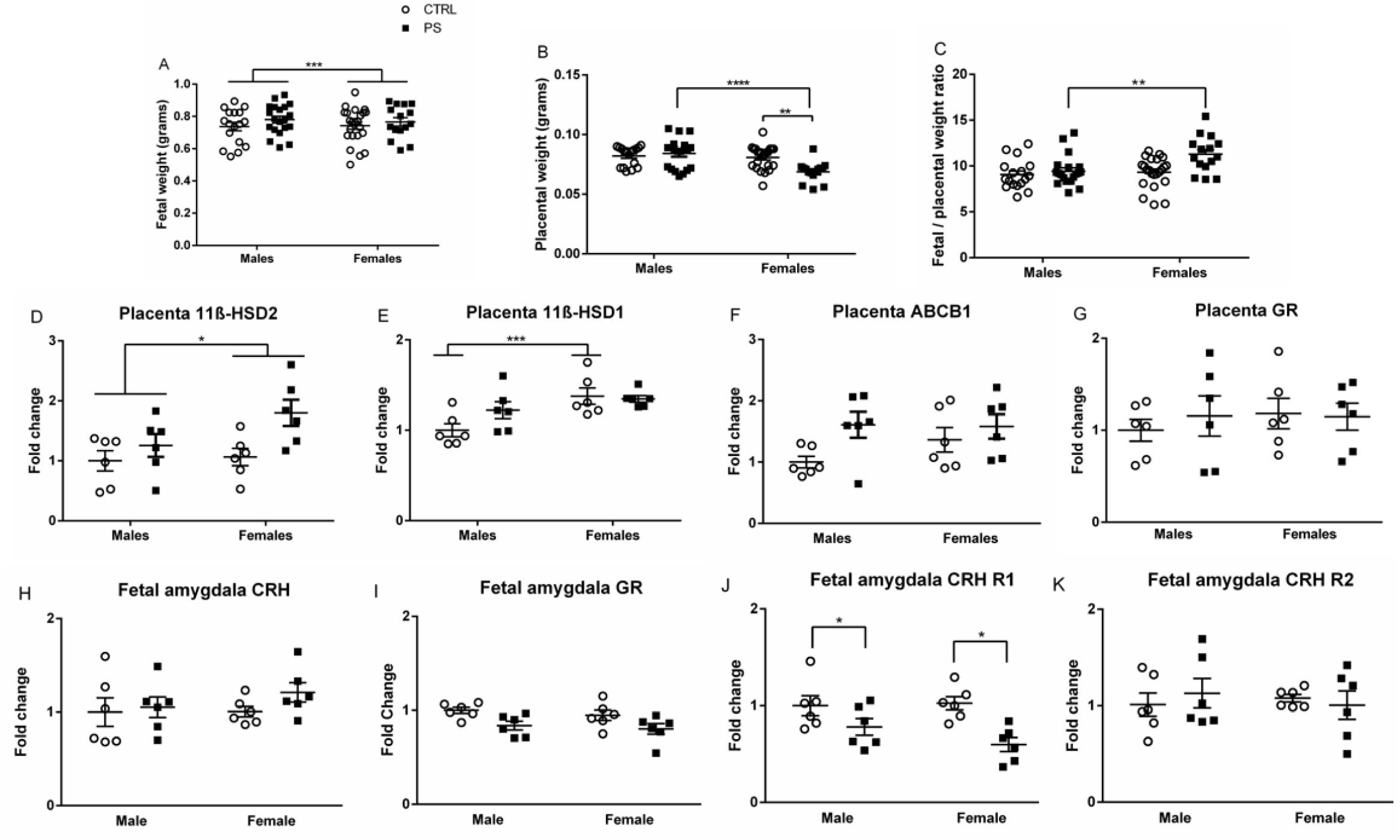
Placental responses to psychosocial stress and effects on fetal brain development. (**A**) Fetal weight (**B**) placental weight and (**C**) fetal to placental weight ratio measurements performed at gestational day 17.5, CTRL = 5 litters, PS = 4 litters. qPCR analysis of placental glucocorticoid barrier components measured at gestational day 17.5 including (**D**) 11ß– HSD2, (**E**) 11ß– HSD1, (**F**) ABCB1, and (**G**) GR, N = 3 litters per group, 2 males and 2 females per litter. qPCR analysis of gene expression changes in molecular regulators of the HPA axis, including (**H**) CRH, (**I**) GR, (**J**) CRH R1, and (**K**) CRH R2, performed in E17.5 fetal amygdalas, N = 3 litters per group, 2 males and 2 females per litter. Data presented as mean ± SEM. \*\**p* < 0.01, \*\*\*\**p* < 0.0001 mixed linear ANOVA with prenatal stress x sex model and litter as a randomized block factor.

We next measured the effects of psychosocial PS on the placental glucocorticoid barrier. There was a significant main effect of sex on 11ß-HSD2 mRNA expression (Figure 4D; F_1,16_= 5.74, P=0.0292), where overall female placentas expressed higher 11ß-HSD2 levels than male placentas at gestational day 17.5. There was a trend for a significant sex x PS interaction (Figure 4D; F_1,16_= 3.58, P=0.0768). PS female placentas had elevated levels of 11ß-HSD2 when compared with CTRL female placentas, although this difference did not reach statistical significance (P=0.0814). For 11ß-HSD1 mRNA, there was a significant main effect of sex (Figure 4E; F_1,16_= 20.87, P=0.0003) and a significant sex x PS interaction (Figure 4E; F_1,16_= 5.27, P=0.0355). CTRL male placentas expressed significantly less 11ß-HSD1 than CTRL female placentas at gestational day 17.5 (P=0.0002), a difference that was no longer present after PS exposure (P=0.1278). Analysis of placenta ABCB1 expression revealed no significant effect of sex (Figure 4F; F_1,16_= 1.68, P=0.2136) or PS (Figure 4F; F_1,4_= 1.70, P=0.2626). As the glucocorticoid receptor (GR) is known to regulate 11ß-HSD2/1 expression, we assessed for changes in placental GR mRNA levels in CTRL and PS mice. There was no sex effect (Figure 4G; F_1,16_= 0.64, P=0.4357) or PS effect (Figure 4G; F_1,4_= 0.06, P=0.8116) for placental GR expression.

To examine possible effects of psychosocial PS on fetal brain development, we investigated whether changes in gene expression could be measured in the amygdala, a key brain region for emotional output, at E17.5. We focused on key molecular regulators of the HPA axis as previous studies have shown these to be altered by PS (Brunton, 2013). We observed a trend towards a significant main effect of PS on CRH mRNA levels (Figure 4H; F_1,4_= 5.39, P=0.0810). CRH levels were elevated in PS fetuses when compared with CTRLs, although this difference was not statistically significant. We also detected a trending effect of PS on amygdalar GR expression (Figure 4I; F_1,4_= 6.84, P=0.0591). PS fetuses exhibited reduced but not statistically significant GR mRNA levels when compared with CTRL fetuses. There was a significant main effect of PS on CRH R1 expression (Figure 4J; F_1,4_= 16.14, P=0.0159). Psychosocial PS exposure resulted in decreased amygdalar CRH R1 mRNA levels in E17.5 fetuses. Analysis of CRH R2 expression revealed no significant effect of PS (Figure 4K; F_1,4_= 0.04, P=0.8449).

These results are summarized in Table 1 and suggest that early alterations in expression of HPA axis regulators within amygdalar neurons may affect neural circuits which lead to the behavioral abnormalities observed in PS offspring.

**Table 1.** Summary of sex-specific placental responses to stress and gene expression changes in fetal amygdala.

| Analysis | N | Female vs. male difference | PS vs. CTRL difference |
| --- | --- | --- | --- |
| Fetal and placental measurements |  |  |  |
| Fetal weight | N=5 litters for CTRL<br>(17 M, 23 F)<br>N= 4 litters for PS<br>(20 M, 15 F) | ↓ F | - |
| Placental weight |  | - | ↓ PS F |
| Fetal/placental weight ratio |  | - | ↑ PS F |
| G17.5 Placenta |  |  |  |
| 11β-HSD2 | N=3 litters per group<br>(2 M, 2 F per litter) | ↑ F | ↑ PS F<br>(Trend)<br>P=0.0814 |
| 11β-HSD1 |  | ↑ F<br>(CTRL F vs. CTRL M only) | Difference no longer after PS |
| ABCB1 |  | - | - |
| GR |  | - | - |
| Fetal E17.5 amygdala |  |  |  |
| CRH | N=3 litters per group<br>(2 M, 2 F per litter) | - | ↑ PS<br>(Trend)<br>P=0.0810 |
| GR |  | - | ↓ PS<br>(Trend)<br>P=0.0591 |
| CRH R1 |  | - | ↓ PS |
| CRH R2 |  | - | - |

### RNA-seq analysis reveals sex-specific alterations in synaptic components and neurotransmitter systems following psychosocial PS

To investigate the molecular mechanisms underlying sex-specific effects of psychosocial PS on behavior and neuroendocrine function, we performed RNA-seq on PN28 amygdalar samples from CTRL and PS male and female offspring, with 4 biological replicates each (sixteen samples in total). Principal component analysis revealed that samples from PS mice were distinguishable from CTRL mice more significantly in male than in female samples (Figure 5A). To examine detailed transcriptomic changes, we performed differential gene expression analysis using four pairwise comparisons: CTRL male vs. PS male, CTRL female vs. PS female, CTRL male vs. CTRL female, and PS male vs. PS female. Differentially expressed genes were determined by FDR < 0.05 and FC > 1.5. Indeed, male mice displayed much larger number of genes differentially expressed after PS exposure than females (Figure 5B).

**Figure 5.**
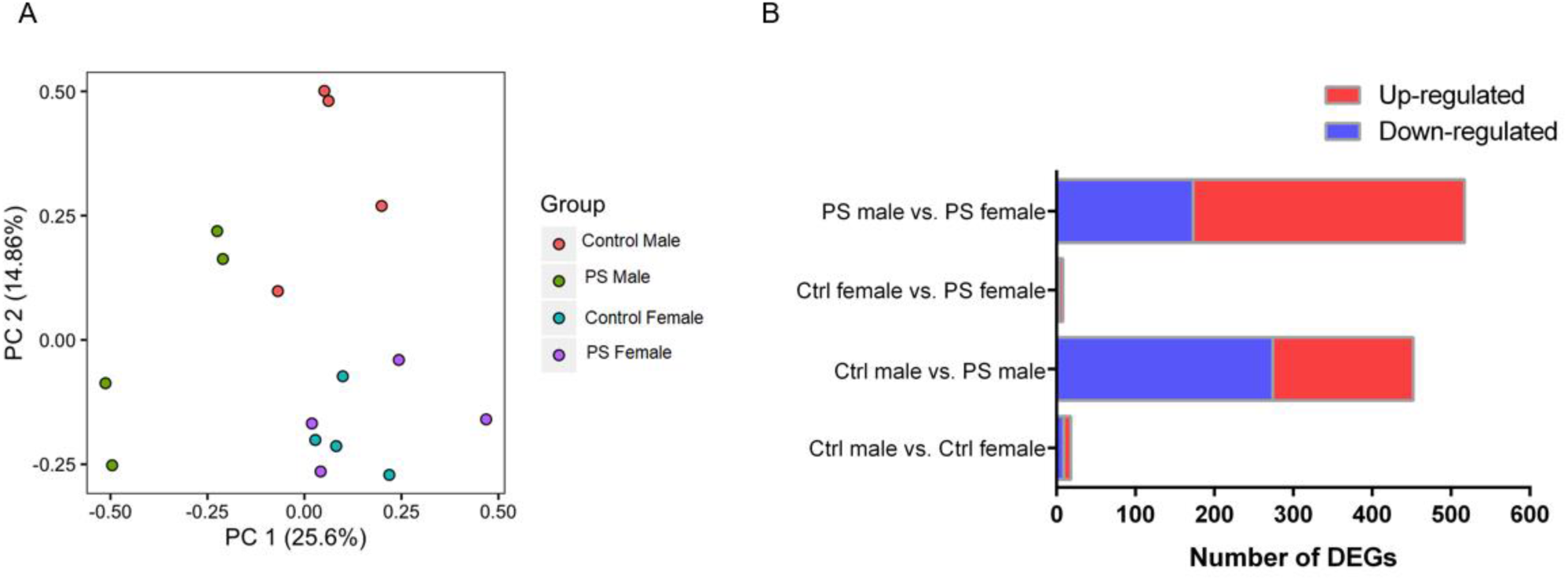
Psychosocial PS results in sexually dimorphic gene expression changes in offspring amygdala. (**A**) Principal component analysis plot. (**B**) Total number of differentially expressed genes for the various pair-wise comparisons made.

In the CTRL male vs. CTRL female amygdala comparison, there were 18 differentially expressed genes (DEGs), with 9 female-specific and 9 male-specific genes (Figure 5 – figure supplement 1A). Transcripts displaying the largest effect were localized on sex chromosomes (*Xist, Eif2s3y, Kdm5d, Ddx3y, Uty*) (Figure 5 – figure supplement 1B), and previously reported to be differentially expressed between the sexes in the hypothalamus (Mozhui et al., 2012). Gene ontology (GO) analysis for the DEGs in this comparison revealed enrichment in genes encoding for calcium binding proteins associated with a wide variety of processes, including inflammation (RAGE receptor binding) and energy metabolism (long chain fatty acid binding) (Figure 5 – figure supplement 1C). Specifically, the S100 genes (*S100a9, S100a8*, and *S100a4*) were expressed in CTRL males when compared with CTRL females suggesting this family of calcium binding proteins may play an important role in modulating amygdalar sexually dimorphic neuronal processes, as has been attributed to other families of calcium binding proteins (Donato et al., 2013; Lephart, 1996).

In the CTRL male vs. PS male comparison, we identified a total of 452 DEGs with 178 up-regulated and 274 down-regulated (Figure 6A). According to the GO analysis, we observed that genes related to regulation of transmembrane transport (including ion channels) and components of the synaptic membrane were significantly down-regulated in PS males (Figure 6 B). The majority were genes associated with potassium channels (including *KCNF1, KCNH4, KCNK9, KCNA3, KCNV1*), genes associated with calcium signaling (*ATP2B4, TRDN, CACNG6, CRACR2A, and CAMK2D*), genes implicated in glutamatergic signaling (*GSG1L, CNIH3, CACNG5, GRIN3A*), and GABAergic signaling (*GABRQ, GABRA3, HAP1*) (Figure 6B). Although GO analysis among the up-regulated genes in PS males did not reveal any significant pathway enrichment (data not shown), several of the significantly up-regulated genes were also associated with glutamatergic signaling (including *GRM2, GRM4, HOMER 3, TCF7L2, GRID2IP*). These results suggest psychosocial PS exposure in males results in alterations in amygdalar synaptic transmission related to glutamate and GABAergic signaling. This may lead to an imbalance between neuronal excitation and inhibition that could be underlying the behavioral abnormalities in PS males observed in our study, as a similar imbalance has been observed in patients with depression or anxiety symptoms (Babaev et al., 2018; Luscher et al., 2011; Murrough et al., 2017). Potassium channelopathy was also noted in PS males which, along with abnormalities in calcium signaling, could imply long-term modifications in membrane excitability and synaptic firing ability.

**Figure 6.**
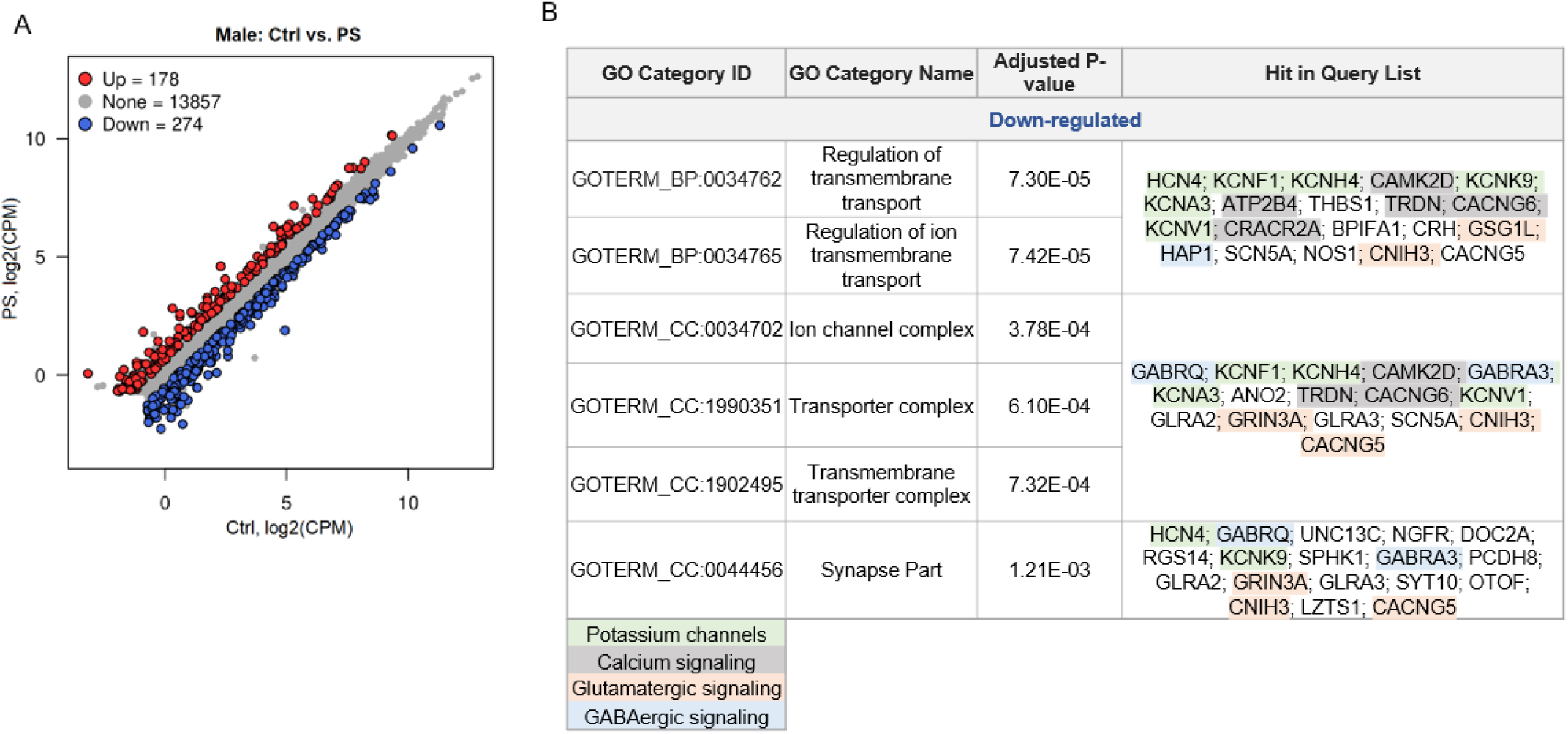
Analysis of DEGs in the amygdala in CTRL males vs. PS males. (**A**) Scatter plot displaying 178 up-regulated and 274 down-regulated genes in PS males when compared with CTRL males. Genes were considered significant with an FDR < 0.05 and FC > 1.5. (**B**) Significantly enriched pathways after gene ontology analysis of down-regulated genes using biological process and cellular component categories. Genes significantly down-regulated in the amygdala of PS males when compared to CTRL males include genes associated with potassium channels, calcium signaling, glutamatergic signaling, and GABAergic signaling.

Interestingly, only a very small subset of genes were differentially expressed in PS females compared with CTRL females. This comparison yielded only a total of 8 DEGs, with 5 up-regulated and 3 down-regulated (Figure 7A). Among the genes displaying the largest effects were *SLC6A5*, involved in glycine neurotransmitter uptake, and *DBH*, which encodes the rate-limiting enzyme for norepinephrine (NE) biosynthesis, which were significantly up-regulated after PS (Figure 7B). Enhanced amygdalar NE activity could be partially underlying the HPA axis hyperactivity phenotype and behavioral abnormalities in females following psychosocial PS.

**Figure 7.**
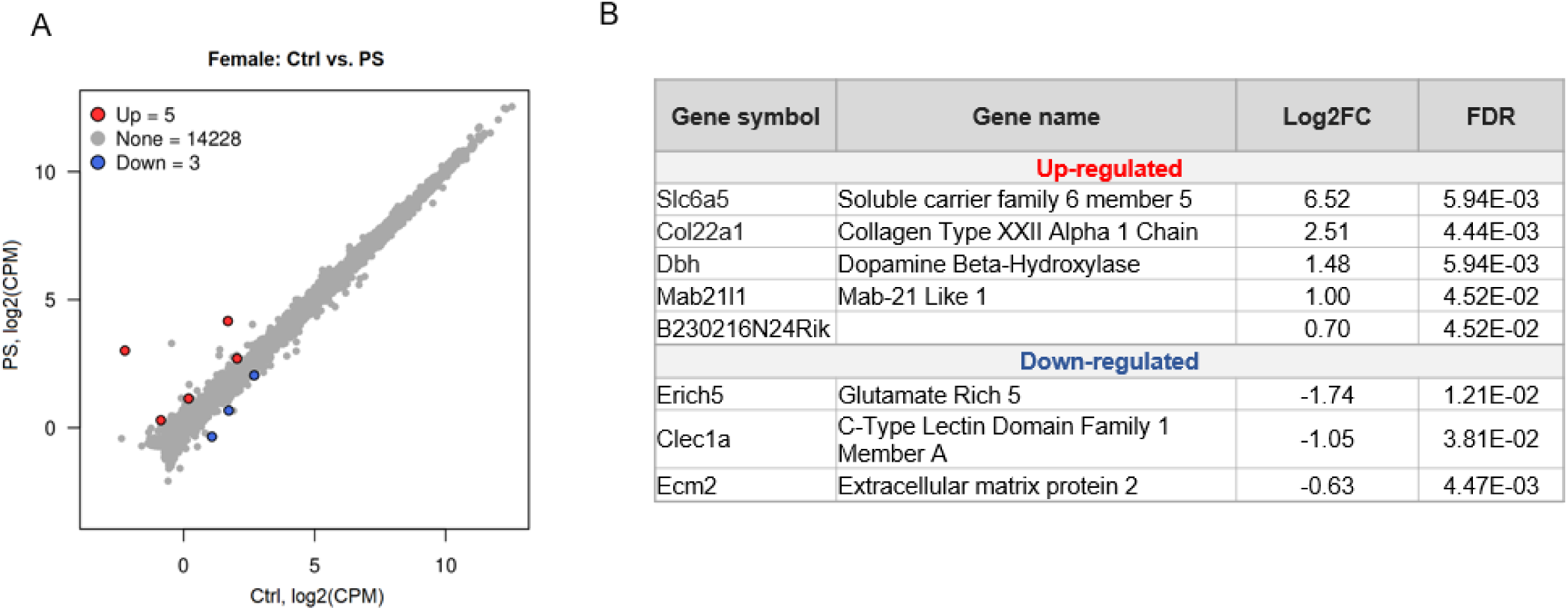
Analysis of DEGs in the amygdala in CTRL females vs. PS females. (**A**) Scatter plot displaying 5 up-regulated and 3 down-regulated genes in PS females when compared with CTRL females. (**B**) Summary of DEGs. Genes were considered significant with an FDR < 0.05 and FC > 1.5.

In the final comparison, PS male vs. PS female, we identified 517 DEGs, with 344 PS female-specific and 177 PS male-specific genes (Figure 7 – figure supplement 1A). To understand how psychosocial PS could have resulted in such distinct patterns of amygdalar gene expression changes in PS males compared with PS females, we performed GO enrichment analysis for the DEGs. Analysis with the biological process category and cellular component category confirmed alterations in synaptic transmission, transmembrane transport, and glutamatergic receptor signaling, as noted above, with genes under these categories significantly expressed in PS females when compared with PS males (Figure 7 – figure supplement 1B). Although GO enrichment analysis among the PS male-specific genes did not reveal any significant pathway enrichment (data not shown), amongst the significantly expressed genes in PS males when compared with PS females were genes associated with steroid dehydrogenase activity (*11ß-HSD1, SRD5A1*). These results suggest differences in steroid metabolism between PS females and PS males could have led to more pronounced gene expression changes in PS males.

### Changes in maternal environment partially rescue sex-specific alterations in genes associated with neurotransmitter systems

Several of the genes associated with synaptic transmission were chosen for qPCR validation. In PS males, we were able to confirm significant alterations in the amygdala of genes encoding proteins involved in glutamatergic (down-regulation of *GSG1L, GRIN3A, CNIH3, CACNG5* and up-regulation *of HOMER 3 and GRM2)* and GABAergic neurotransmission (down-regulation of *GABRA3A*) (Figure 8A-B). Down-regulation of genes associated with ion channel complexes was also confirmed (*KNCH4, KCNA3, CACNG6*) (Figure 8C). *GABRQ* was the only gene chosen for validation via qPCR which exhibited a trend towards down-regulation after PS in males (P=0.0837), a direction of change consistent with RNA-sequencing results, but that did not reach statistical significance (Figure 8A). In females, there was a significant up-regulation in DBH after PS, and a trend towards increased levels of *SLC6A5* (FIGURE 8D; P=0.0899).

**Figure 8.**
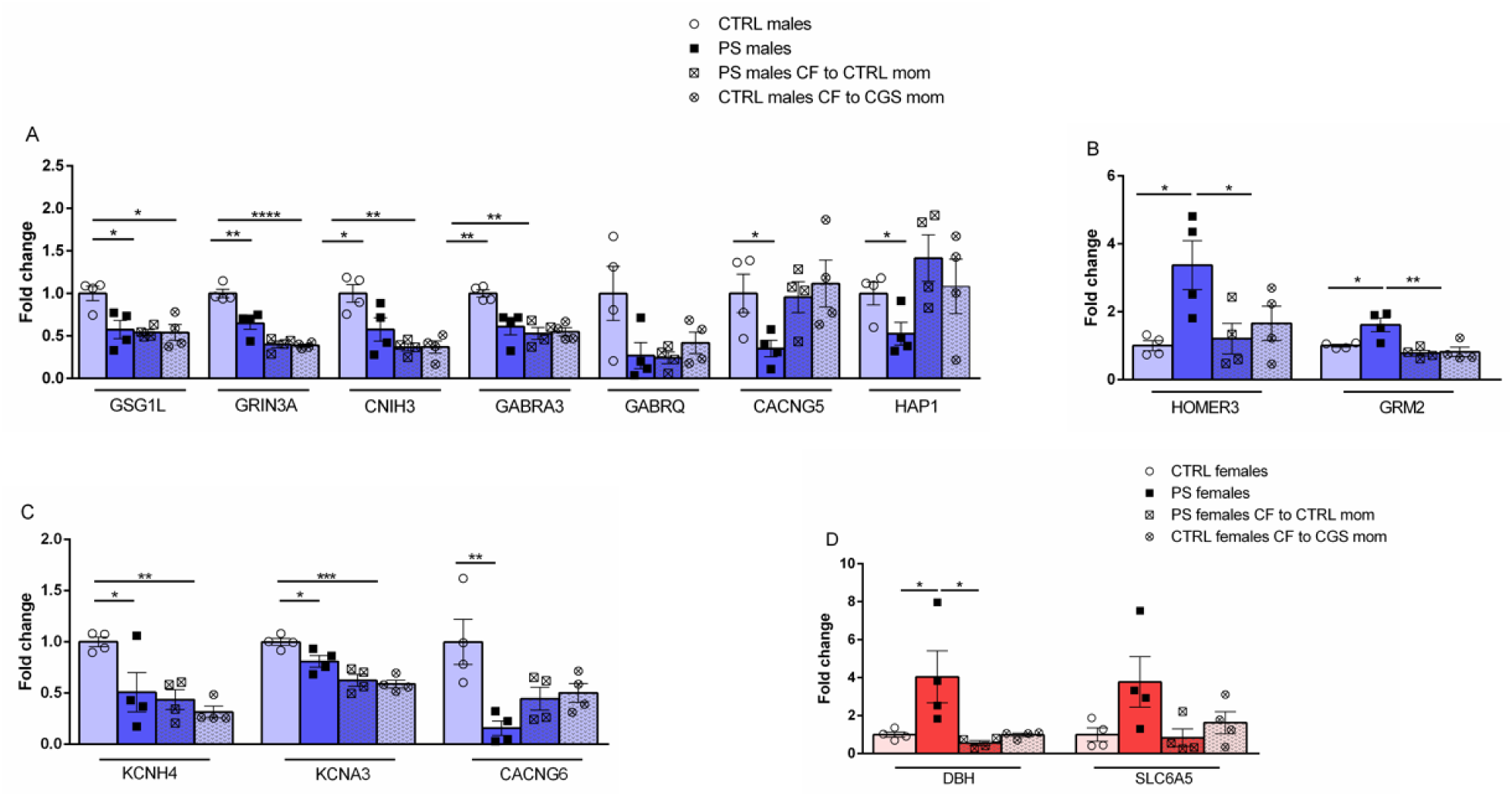
Validation of RNA-Seq results through qPCR and effects of CF on gene expression. (**A**) Validation of RNA-seq results for a subset of genes confirms down-regulation in genes associated with glutamatergic and GABAergic neurotransmitter systems in PS males that are not reversed by changes in postnatal maternal environment. (**B**) Reversal of genes associated with metabotropic glutamate neurotransmission are observed in PS males following CF. (**C**) qPCR data also confirms down-regulation of genes encoding ion transporter complexes in PS males which are not reversed by alterations in maternal environment. (**D**) Validation of RNA-seq results in PS females confirms up-regulation in DBH and a trend towards increased levels in SLC6A5 transporter. CF induces normalization of alterations in DBH in PS females, associated with NE neurotransmission. N = 4 per group. Data presented as mean ± SEM. *p<.05, \*\**p* < 0.01, ***<p<.001, \*\*\*\**p* < 0.0001, unpaired 2-tailed *t* test for differences between Ctrl and PS offspring, two-way ANOVA followed by Tukey’s post hoc test for CF gene expression analysis with PS x CF model.

To differentiate the contribution of in-utero stress from maternal environment on gene transcription, we measured how CF altered PS-induced gene expression changes. We observed that most of the genes down-regulated in PS males were also down-regulated in CTRL males CF to CGS moms, suggesting both in-utero stress and alterations in maternal care are contributing to transcriptional changes noted in our study (Figure 8A and 8C). Importantly, in PS males CF to CTRL moms, we observed a partial rescue of alterations in genes associated with glutamatergic signaling. We find expression levels of *HOMER3* (Figure 8 B; F(1,12) = 7.990; P = 0.0153 PS × CF interaction; post-hoc P < 0.05) and *GRM2* (Figure 8 B; F(1,12) = 6.178; P = 0.0287 PS × CF interaction; post-hoc P < 0.01) are normalized following CF, and note a trend towards normalization of *CACNG5* and *HAP1* (Figure 8A), suggesting a partial rescue in metabotropic glutamatergic neurotransmission could underlie the reversal of anxiety-related behaviors in PS males CF to CTRL moms. In PS females, we observe expression of *DBH* is normalized following CF to CTRL moms (Figure 8D; F(1,12) = 6.235; P = 0.0281 PS × CF interaction; post-hoc P < 0.05), suggesting a partial rescue in NE neurotransmission could underlie the reversal of anxiogenic behaviors in PS females CF to CTRL moms.

Our results are summarized in Figure 8-Supplement 1.

## DISCUSSION

In our study, we found that psychosocial PS exposure results in the emergence of anxiety-like behaviors, an increased state of alertness displayed as decreased amount of time spent swimming in the FST, and anhedonia in male and female offspring. Neuroendocrine abnormalities, evidenced by acute stress induced HPA axis hyperactivity, are only observed in female offspring after PS. We further demonstrate these behavioral and neuroendocrine abnormalities are differentially modulated by changes in maternal care in the postnatal environment, with only anxiety-related behaviors able to be rescued by CF to CTRL mothers. In addition, we find evidence that these abnormalities are preceded by sex-specific placental responses to PS and gene expression changes in the fetal amygdala, suggesting programming of brain development leading to an increased susceptibility to emotional disturbances. Molecular analysis further revealed phenotypic changes in PS offspring to be associated with sex-specific disturbances in genes associated with synaptic transmission. PS males display alterations in genes encoding ion channels, transporter complexes, and components of the synaptic membrane, including genes associated with glutamatergic and GABAergic signaling. In contrast, PS females displayed notably fewer abnormalities in gene expression as compared to the CTRLs. However, *DBH* which regulates NE synthesis was significantly upregulated and NE is well known to regulate numerous behaviors mediated by the amygdala (Levy & Tasker, 2012; Niwa et al., 2011). Lastly, CF of PS offspring to CTRL mothers partially normalizes changes in gene expression related to glutamatergic and NE signaling, which may contribute to the reversal of anxiety-like behaviors.

Our data indicate exposure to psychosocial PS leads to anxiety-like behaviors and anhedonia in male and female offspring. In contrast, only PS female offspring exhibit HPA axis dysregulation. Consistent with our data, an abundance of studies have shown PS exposure results in depressive behaviors and anhedonia in offspring of both sexes (Weinstock, 2016). However, previous studies reporting the effects of PS on anxiety-related behaviors and neuroendocrine dysfunction are not always consistent. PS paradigms with a psychosocial component have demonstrated anxiety-like behaviors and HPA axis hyperactivity in male offspring (Abe et al., 2007; Brunton & Russell, 2010; Mueller & Bale, 2008), with males being particularly susceptible to effects of PS experienced during early gestation (Mueller & Bale, 2008). Interestingly, increased anxiety and stress-induced HPA axis hyperactivation are reported in PS female offspring if stress paradigms with a psychosocial component are applied during middle to late gestation (Bosch et al., 2007; Soares-Cunha et al., 2018). These results suggest sex-specific windows of vulnerability to psychosocial PS exposure, with females becoming susceptible to HPA axis dysfunction and the development of anxiety during mid to later stages of fetal development. These sex-dependent effects of PS could partially be explained by developmental timing of relevant processes, including a greater increase in limbic GR expression in female brains during later in-utero life (Owen & Matthews, 2003). Of note, data from human studies support the notion that the HPA axis in females might be more sensitive to programming as several studies have shown females exposed to various prenatal insults exhibit increased HPA axis reactivity compared with males when challenged with a maternal separation event or the Tier Social Stress Test (Carpenter et al., 2017).

Our findings further demonstrate sex-specific alterations in placental responses to PS. The placenta plays a critical role in fetal nutrition during pregnancy, orchestrating the active transport of nutrients and metabolic waste from maternal to fetal circulation (Gaccioli & Lager, 2016). Here, we show that placentas from PS females weigh significantly less than placentas from CTRL females. Despite this reduction in placental weight, PS female fetuses weight is maintained, and PS females display an increased fetal to placental weight ratio, suggesting enhanced placental efficiency in supporting fetal growth. These results further suggest placentas from females exhibit a higher capacity to adapt to suboptimal intrauterine conditions through morphological or functional changes (Sferruzzi-Perri & Camm, 2016). Consistent with this notion, placentas from PS females exhibit a trend towards increased 11ß-HSD2 mRNA, suggesting a compensatory response to metabolize excess glucocorticoids. Activity of placental 11ß-HSD2 also appears to be sex-linked in humans, with more adaptive responses found in females (Stark et al., 2009). Several studies have shown distinct sex differences in epigenetic regulators and methylation patterns of placental tissues, providing a plausible mechanism by which females may adopt placental adaptive responses to environmental insults (Bale, 2016; Gallou-Kabani et al., 2010; Nugent et al., 2018). It remains to be assessed in our model if sex specific effects of PS on histone modifications or microRNA expression in the placenta could account for differences in placental responses we observed.

Interestingly, previous studies have shown 11ß– HSD2 expression to be affected by PS, in a temporal and sex-specific manner. In mice, exposure to chronic variable stress during the first week of gestation results in a reduction in placental 11ß– HSD2 in females and a trend toward increased expression in males (Pankevich et al., 2009). Chronic restraint stress or dietary protein restriction during mid to late gestation in rats results in decreased 11ß– HSD2 expression in both sexes (Belkacemi et al., 2011; Jensen Peña et al., 2012; Lesage et al., 2001), and increased placental 11ß– HSD2 is observed in low anxiety but not high anxiety bred rats after exposure to chronic social defeat (Lucassen et al., 2009). Here, we show 11ß– HSD2 expression is modulated differently by sex and psychosocial stress, with a trend toward increased levels in females observed in late gestation. We did not observe any significant effect of PS on 11ß– HSD1 or ABCB1. Expression levels of GR, known to regulate transcription of 11ß– HSD2, were also not affected by PS. These results suggest other mechanisms, such as crosstalk between estrogen receptor and protein kinase A signaling pathways, which are associated with transcriptional regulation of hydroxysteroid dehydrogenases (Chapman et al., 2013; Guan et al., 2013), might be underlying the moderate sex-specific increase in 11ß– HSD2 noted in our study.

Importantly, despite the placental responses observed, both PS male and female fetuses exhibited changes in amygdalar gene expression. Psychosocial PS exposure was associated with decreased CRHR1 expression in both sexes as well as trends towards up-regulation of CRH mRNA levels and down-regulation in GR expression. These results indicate the placental 11ß– HSD2 barrier is being overcome by elevated levels of glucocorticoids, or reduction in enzymatic activity, resulting in fetal overexposure to CORT and alterations in brain developmental trajectories. An alternative, or additional contributor, might be glucocorticoids of fetal origin as fetal HPA axis has been shown to be activated by maternal adversity during late gestation (Fujioka et al., 2003; Ohkawa et al., 1991). Similarly, other stress-related amine hormones, such as catecholamines, known to regulate various placental processes and influence brain development, might be involved (Bronson & Bale, 2016).

The amygdala undergoes dramatic structural and functional modifications as a result of stress (Schulkin, 2017; Zhang et al., 2018). Here, we examined amygdalar transcriptional responses following psychosocial PS. Enrichment analysis revealed PS males exhibited alterations in genes involved in synaptic transmission, including down-regulation of voltage gated potassium channels and genes involved in calcium signaling. In addition, we detected alterations in transcripts encoding synaptic proteins involved in glutamatergic and GABAergic neurotransmitter systems. Our findings are consistent with other studies examining effects of prenatal insults on amygdalar gene expression which have observed changes in ion transporter complexes, GABA_A_ receptor subunits, as well as ionotropic and metabotropic glutamate receptors (Barrett et al., 2017; Ehrlich et al., 2015; Laloux et al., 2012). Our data, however, identify a number of novel PS responsive transcripts, including genes encoding regulatory proteins known to modulate AMPA receptor trafficking and channel kinetics, such as *GSG1L, CACNG5*, and *CNIH3* (Bissen et al., 2019; Gu et al., 2016), genes encoding proteins involved in postsynaptic stabilization of metabotropic glutamate receptors, such as *HOMER3*, and GABA_A_ receptor trafficking partner *HAP1* (Figure 9) (Mandal et al., 2011; Twelvetrees et al., 2019). Proper balance of excitatory and inhibitory neurotransmission is crucial for preventing amygdala hyperactivity, a neurological feature often noted in patients suffering from depression and anxiety disorders (Tzanoulinou et al., 2014; Zhang et al., 2018). The complex pattern of changes in genes associated with synaptic transmission detected in our study could be affecting the balance of excitatory and inhibitory transmission within the amygdala and resulting in the behavioral abnormalities observed in PS males.

**Figure 9.**
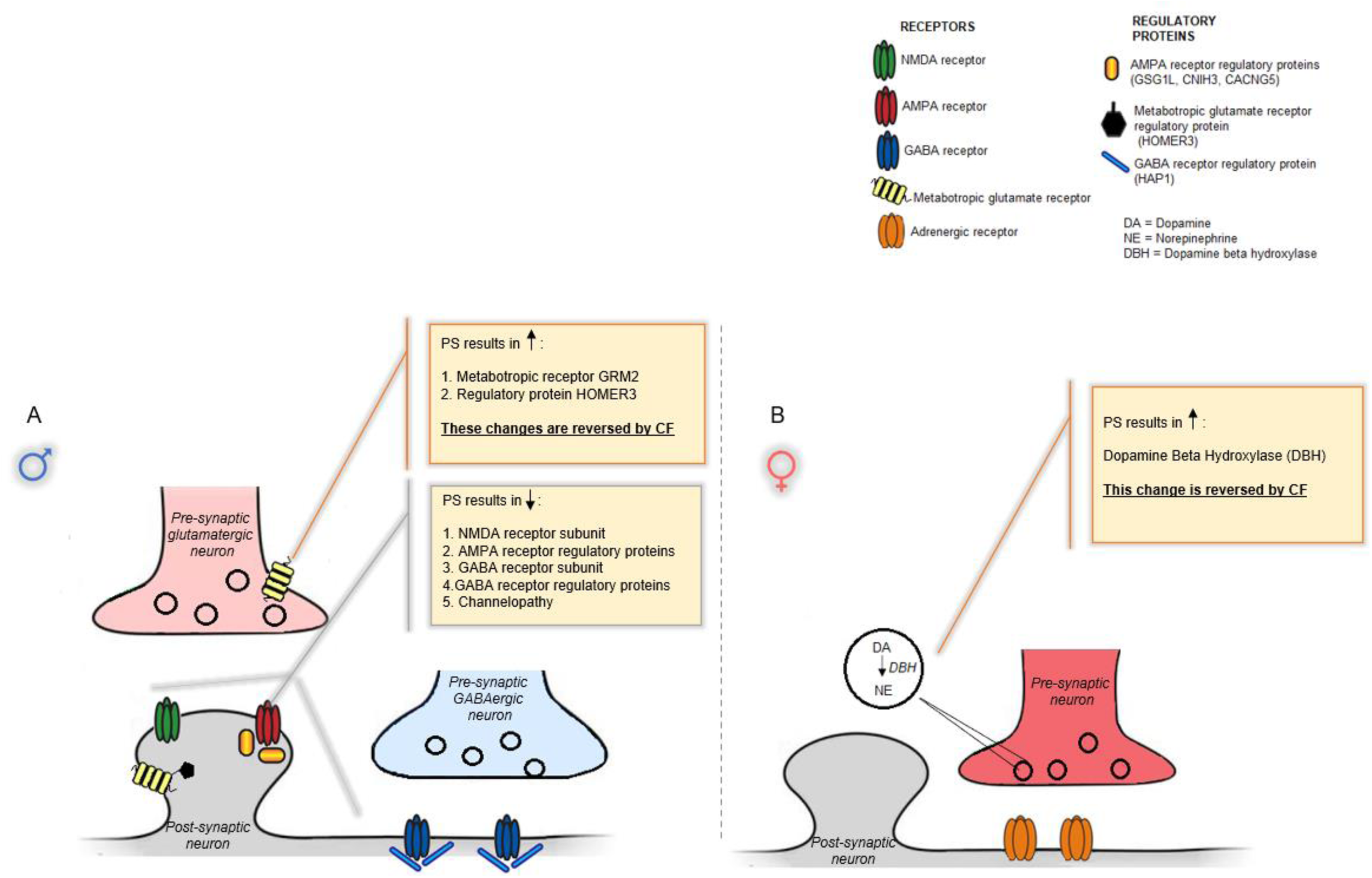
Model summarizing gene expression changes in PS offspring and normalizations induced by CF. (**A**) In males, psychosocial PS results in down-regulation genes encoding proteins associated with glutamatergic signaling, GABAergic signaling, and ion channel transporter complexes. These transcriptional changes associated with synaptic transmission are likely resulting in an imbalance in excitatory and inhibitory transmission, resulting in the behavioral abnormalities observed in PS males. CF to CTRL mothers normalizes gene expression changes associated with metabotropic glutamate receptor signaling and reverses anxiety-related behaviors. (**B**) In females, PS results in an up-regulation of DBH, which is likely to increase noradrenergic tone and partially underlie HPA axis hyperactivity and increased anxiety noted in PS females. CF normalizes DBH expression and reverses anxiety-related behaviors in PS females.

Unexpectedly, only minor changes were observed in PS females, including increased expression of dopamine beta hydroxylase (*DBH*), a gene associated with NE synthesis and this suggesting increased NE signaling (Figure 9). Stress induced increases in *DBH*, which catalyzes the conversion from dopamine to NE (Fan et al., 2013), can result in increased noradrenergic tone and partially underlie the enhanced stress sensitivity and increased anxiety noted in PS females (Levy & Tasker, 2012; Niwa et al., 2011). Of note, a significant down-regulation of genes associated with steroid metabolism, including 11ß-HSD1, were noted in PS females when compared with PS males. Decreased 11ß-HSD1 expression in the amygdala of PS females could serve as a protective factor, exposing this brain region to lower CORT levels than PS males, thus partially explaining the gender specificity of phenotypes observed. Alternatively, the lack of robust gene expression changes observed in PS females could suggest alterations not detectable at the transcriptional level, such as protein post-translational modifications, could be mediating female neuronal responses to stress in the amygdala. Other brain regions that exhibit a robust sexually dimorphic response to stress, such as the bed nucleus stria terminalis (Carvalho-Netto et al., 2011; Herman & Tasker, 2016), could also be affected.

A primary objective of our study was to dissociate in-utero from postnatal maternal environmental effects. In line with other studies, our data indicate that in-utero stress contributes to the emergence of anxiety-related behaviors, anhedonia, and HPA axis dysfunction (Brunton, 2013; Weinstock, 2008, 2017). Interestingly, we find these abnormalities are differentially affected by changes in maternal care. Anxiety-like behavior exhibited by PS offspring are rescued by CF to CTRL mothers, suggesting alterations in neural circuits underlying this phenotype are ameliorated by changes in quality of maternal care received. Consistent with this notion, we observed normalization of amygdalar *HOMER3* and metabotropic glutamate receptor *GRM2* expression following CF of PS males with CTRL dams as well as rescue of *DBH* expression in PS females CF to CTRL moms (Figure 9). These results suggest CF-induced changes in metabotropic glutamate signaling and NE signaling could be partially restoring the balance in excitatory and inhibitory transmission within the amygdala of PS offspring, thus ameliorating anxiogenic behaviors observed. Previous studies manipulating quality of maternal care by neonatal handling have shown the reversal of both anxiogenic behavior and HPA axis hyperactivity following PS (Maccari et al., 1995; Vallée et al., 1997; Weinstock, 2015). In our study, cross-fostering was not effective in rescuing the enhanced HPA axis drive and increased state of vigilance noted in PS offspring, suggesting the influence of psychosocial PS on these phenotypes differs from the effect of other types of prenatal insults, with psychosocial PS being associated with more persistent deleterious alterations. Lastly, the emergence of anhedonia in PS offspring observed in our study seemed to be mediated both by in-utero stress and abnormalities in maternal care, and was not ameliorated with CF. Gene expression alterations not normalized by CF might be contributing to these phenotypes in PS males, including down-regulation of several regulatory proteins associated with AMPA receptor trafficking, ion channel complexes, and GABA_A_ receptor subunit *GABRA3*.

In conclusion, we find psychosocial PS results in behavioral and sex-specific neuroendocrine abnormalities that are differentially modulated by changes in maternal care. These abnormalities are associated with a complex pattern of gene expression changes in the amygdala indicating dysfunctions in synaptic transmission and neurotransmitter systems with pronounced sex differences. In addition, we find CF triggers changes in genes associated with glutamatergic and NE neurotransmission that are associated with reversal of anxiety-related behaviors. Future experiments are needed to investigate cellular and electrophysiological changes in amygdalar structures that may result from the molecular alterations identified in our study. Studies aimed at interrogating the role specific genes altered by CF play in the rescue of anxiogenic behaviors could provide opportunities for the development of novel, clinically relevant therapeutic strategies.

## MATERIALS AND METHODS

### Animals

C57BL6/J mice were obtained from Jackson Laboratory. Mice were housed on a 14-hour/10-hour light-dark cycle (lights on at 6:00 A.M.) with access to water and chow *ad libitum*. After being allowed to habituate to the animal facility for at least 2 weeks, virgin female mice between 3 and 6 months of age were set up for timed-mating at 1800h and separated the following morning at 0800h. Simple randomization was used to divide female mice with a copulatory plug, which was denoted as 0.5 days post-coitum, into two experimental groups [CTRL and stressed (CGS)], and housed in groups of four. All mouse experiments were in accordance with the guidelines of the National Institutes of Health and were approved by the Cincinnati Children’s Medical Center Animal Care and Use Committee under IACUC protocol number 2017-0051.

### Psychosocial prenatal stress

Chronic psychosocial stress paradigm during pregnancy (CGS) was conducted as previously described (Zoubovsky et al., 2020). Briefly, from gestational day 6.5 to 16.5, mice assigned to the stress group were exposed to variable psychosocial insults 2 times per day, 2 h each, and an overnight stressor. Stressors included rat odor exposure, foreign object exposure, 30° cage tilt, bedding removal, frequent bedding changes, overnight lights on, overnight wet bedding, and overnight cage mate change. Control mice were not disturbed.

### Cross-fostering (CF)

On the day of birth (postnatal day 0, PN0), litters were culled to 3 male and 3 female pups per dam to ensure similar conditions across all dams. For CF litters, pups were switched with another litter within 24h of birth, so as to generate four groups for our studies: CTRL offspring, PS offspring, PS offspring CF to CTRL mom, CTRL offspring CF to CGS mom.

### Offspring Behavioral Assessment

Behavioral tests were conducted from least to most stressful in offspring when they reached PN28. Three separate cohorts were used for behavioral testing. The first cohort underwent testing for anxiety-related behaviors using LD, followed by OFT to assess for changes in locomotor activity, SI to measure alterations in sociability, and FC to quantify deficits in associative learning at the Cincinnati Children’s Research Foundation’s Animal Behavior Core. A second cohort was used to assess for depressive-like behaviors using the FST. A third cohort was used to measure changes in anhedonia using the SPT. To control for litter effects, one to two males or one to two females per litter selected by simple randomization were tested. Statistically, litter was a randomized block factor during analysis. Experimenters were blinded to group membership.

#### Light dark transition box (LD)

The LD was performed as previously described (Amos-Kroohs et al., 2013). Briefly, offspring were placed inside the apparatus used for OFT which was modified with a black acrylic insert that divided the chamber into two sides, one dark and one light, each measuring (20.5 cm × 41 cm) with a 7 × 7 cm opening between them. Mice were placed on the lighted side and the amount of time spent in each side of the apparatus as well as number of crossings was recorded over a 10-min period.

#### Open field test (OFT)

The OFT was conducted as previously described (Kuerbitz et al., 2018). Briefly, mice were placed inside an activity chamber measuring (41 cm x 41 cm x 38 cm) (San Diego Instruments, San Diego) with 16 photobeams spaced 2.5 cm apart in the x and y planes. Mice were tested for 1h and locomotor activity was analyzed in 5-min intervals.

#### Social interaction assay (SI)

SI was performed as previously described (Amos-Kroohs et al., 2016) with minor modifications. A three-chamber clear acrylic apparatus divided in three compartments was used. Two larger compartments (30 × 70 cm each) and a smaller central compartment (8.5 × 70 cm) were connected by openings in the center of each partition (7.5 × 5 cm). The larger compartments contained a small circular confinement cage each (measuring 15 cm in diameter). The test consisted of three parts conducted on a single day. Mice were first placed in the center compartment without access to the larger chambers and allowed to explore for 5 min. Next, mice were allowed to explore all three chambers of the apparatus for 5 min. Finally, in the testing phase, a stranger mouse was introduced (an unfamiliar conspecific of the same strain and sex) into one of the two confinement cages and the other confinement cage remained empty. Mice were allowed to explore freely for 5 min and movement was tracked using ANYmaze software (Stoelting, Wood Dale, IL). Amount of time spent interacting with the stranger mouse was used to quantify the degree of social interaction.

#### Fear conditioning test (FC)

The FC assay which consisted of CS/US training and contextual and auditory cued components for fear conditioning in order to assess associative learning was performed as previously described (Vorhees et al., 2015). The apparatus contained a grid floor connected to a scrambled foot shock device mounted inside a sound-attenuated chamber (San Diego Instruments, San Diego, CA). On day one, mice were placed inside the apparatus for the training phase where they received three tone-footshock pairings spaced 180 s apart (tone: 82 dB, 2 kHz, 30 s duration; shock 1 s, 0.3 mA) near the end of a 10 min period. On day two, for contextual fear testing, mice were placed back in the chamber for 6 min without auditory cues. On day three, for cued fear testing, the grid floor inside the chamber was replaced with a different floor with a black hexagonal insert and mice were placed in the chamber for 3 min with no tone present followed by 3 min of tone. Freezing behavior was quantified on day two and three.

#### Forced swim test (FST)

The FST was performed as previously described (Amos-Kroohs et al., 2016). Briefly, mice were placed in a 2 L beaker with 1.5 L of water acclimatized to room temperature (25°C). On day 1, the habituation session, mice were placed in the beaker for 15-min. The testing session occurred the following day. Mice were allowed to swim for 5-min and duration of immobility as well as frequency of immobility episodes were recorded. Immobility was defined as lack of all motion except the minimal movement required to keep the mouse afloat.

#### Sucrose preference test (SPT)

The SPT was conducted as previously described (Zoubovsky et al., 2020). Briefly, mice were single housed with *ad libitum* chow, and given free access to one 100 ml graduated bottle containing tap water and another 100 ml bottle containing 4% sucrose for 6 days. The position of the bottles was interchanged daily to reduce side-bias. Water and sucrose consumption (ml) were measured and preference was calculated using the average of the measurements from the last 4 days with the following formula: % preference = [(sucrose consumption/sucrose + water consumption) × 100].

### Serum corticosterone (CORT) measurements

In a separate cohort of mice, submandibular bleeds were performed at PN28 at circadian nadir, peak, and immediately following a 15-min swim in water acclimatized to room temperature (25°C). The blood was collected in serum separator tubes, centrifuged at 21,130 x g for 6-min, and serum was removed and stored at - 20°C. Serum CORT measurements were performed by ELISA per manufacturer’s protocols (Arbor Assay, Ann Arbor, MI).

### Mouse tissue collection, amygdala microdissections, and RNA isolation

Pregnant female mice were euthanized on gestational day 17.5. Following laparotomy, the placentas and corresponding fetuses were collected and weighed. Tail tissue samples were used for genotyping to identify sex of individual fetuses with primers specific for SRY (5’-GAGTACA-GGTGTGCAGCTCTA-3’ and 5’-CAGCCCTACAGCCACATGAT-3’) as previously described (Mueller & Bale, 2008). From a separate group of mice, placentas were harvested and immediately flash frozen in liquid nitrogen. Brains from corresponding fetuses were collected and the ventrolateral portions of caudal telencephalic gross sections containing the amygdala were microdissected under a dissecting microscope and stored in RNA*later* (Thermo Fisher Scientific, Waltham, MA) while tail tissue samples were used for sex genotyping. From each litter, two males and two females were selected by simple randomization for RNA isolation. Tissues were homogenized using stainless steel beads in a TissueLyser II apparatus (Qiagen, Hilden, Germany). Placental RNA was purified using the RNeasy Mini Kit (Qiagen) per manufacturer’s protocol. RNA from amygdalar dissections of fetal brain was purified using RNeasy Micro Kit (Qiagen) following manufacturer’s instructions. For PN28 offspring, mice were sacrificed via cervical dislocations and brains were harvested, immediately frozen on dry ice, and stored at −80°C. Frozen brains were then submerged in RNA*later*-ICE (Thermo Fisher Scientific) and allowed to thaw overnight at −20°C per manufacturer’s protocol. Amygdalar dissections were performed as previously described (Zapala et al., 2005). Briefly, two coronal cuts were made at −1 mm and −2.75 mm with respect to bregma and the amygdala was microdissected from these slices under a dissecting microscope following delineations from the mouse brain atlas to ensure that there was no contamination from surrounding tissue. RNA from amygdalar dissections of PN28 brain was purified using RNeasy Micro Kit (Qiagen) according to manufacturer’s instructions.

### quantitative PCR (qPCR)

RNA was converted to cDNA using the Quantitect Reverse Transcriptase Kit (Qiagen) according to manufacturer’s protocols and stored at −20°C. qPCR was performed using Taqman system with Taqman Gene Expression Master Mix (ThermoFisher Scientific) on each cDNA sample. Specific probes were used to quantify murine targets of interest (ThermoFisher Scientific) and GAPDH (Mm99999915_g1, ThermoFisher Scientific) was used as endogenous control. 50 ng of placental cDNA template and 12.5 ng of fetal or PN28 amygdala cDNA template were used per well, and samples were run in duplicates. qPCR reactions were run on an Applied Biosystems StepOnePlus Real-Time PCR instrument (Applied Biosystems, Foster City, CA) and fold induction were calculated using the ΔΔCt method, normalizing experimental groups to the average of a relevant control group.

### RNA-sequencing

RNA isolated from amygdalar microdissections from four control and four PS offspring (per sex) from different litters at PN28 was used for directional RNA-seq performed by the Genomics,, Epigenomics, and Sequencing Core at the University of Cincinnati following previously published methods (Rapp et al., 2020; Walsh et al., 2019). Briefly, RNA quality was determined by Bioanalyzer (Agilent, Santa Clara, CA). PolyA RNA was isolated using the NEBNext Poly(A) mRNA Magnetic Isolation Module (New England BioLabs, Ipswich, MA) with a total of 1 µg of good quality total RNA as input and enriched using the SMARTer Apollo NGS library prep system (Takara Bio USA, Mountain View, CA). Sequencing libraries were prepared using the NEBNext Ultr II Directional RNA Library Prep Kit for Illumina (New England Biolabs). After indexing, enrichment through 8 cycles of PCR, and passing initial quality control metrics, individually indexed and compatible libraries were proportionally pooled and sequenced using Nextseq 550 sequencer (Illumina, San Diego, CA). The sequencing setting of single read 1 × 85 bp to generate ∼50 M reads per sample was used.

### RNA-sequencing analysis

RNA-seq reads were aligned to mouse genome, mm10, using STAR aligner (Dobin et al., 2013). Raw reads counts aligned to each genes were measured using FeatureCounts (Liao et al., 2014). Differential gene expression analysis was done using RUVseq (Risso, 2014) and EdgeR (McCarthy et al., 2012). Specifically, we introduced two factors (k=1) of unwanted variation and estimated it using RUVs in RUVseq, which were incorporated into the model matrix for edgeR. Genes were defined as “expressed” if it displays > 0.5 CPM in at least one condition. Genes with fold-change > 1.5 and FDR < 0.05 were selected as differential genes for gene ontology analysis using EnrichR (Kuleshov et al., 2016). Top significant GO terms were selected for presentation.

### Statistics

Data were analyzed by mixed linear factorial ANOVA with degrees of freedom calculated using the Kenward-Roger method (Proc Mixed, SAS version 9.4, SAS Institute, Cary, NC, USA) or two-way ANOVA test followed by Tukey’s post hoc test (Prism 7.0c software; GraphPad Software, Inc., San Diego, CA, USA) as indicated in figure legends. For behavioral and neuroendocrine assessments mixed linear ANOVA with PS x sex x CF model was used with litter as a randomized block factor. For placental and fetal weights, and placental and fetal amygdala gene expression measurements, mixed linear ANOVA with PS x sex model and litter as a randomized block factor was used. For qPCR validation of RNA seq data analysis was performed by Student’s *t* test. Effects of cross-fostering on amygdalar gene expression was analyzed by two-way ANOVA for PS and CF to determine main effects followed by Tukey’s post hoc test. P≤0.05 was considered significant. The n represents either offspring or litter numbers as indicated in figure legends. Results are reported as mean ± standard error of the mean (s.e.m.).

## ACKNOWLEDGEMENTS

This work was supported by the National Institute of General Medical Sciences T32 GM063483-14 grant, Cincinnati Children’s Hospital Research Foundation, and the University of Cincinnati Office of the Vice President for Research – URC Graduate Student Stipend and Research Cost Program for Faculty—Student Collaboration award to SPZ. Additionally, the authors would like to thank the Genomics, Epigenomics, and Sequencing Core at the University of Cincinnati for their help in generating the transcriptomics data for this project.

