## Supplemental figures for "Prenatal psychosocial stress-induced behavioral and neuroendocrine abnormalities are associated with sex-specific alterations in synaptic transmission and differentially modulated by maternal environment"

**FIGURE 2 – FIGURE SUPPLEMENT 1**

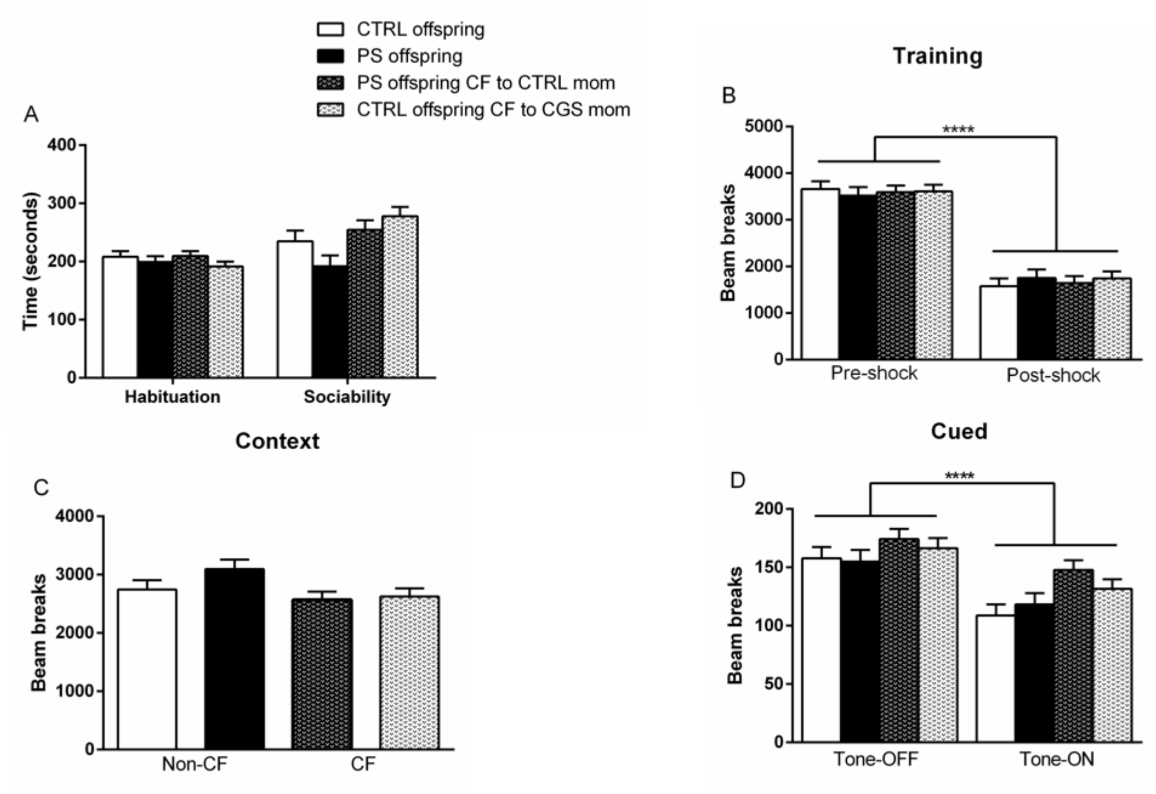


**Figure 2 – figure supplement 1. Psychosocial PS has no effect on sociability or associative learning.** (**A**) Time spent exploring chamber in habituation phase and in testing phase with an unfamiliar conspecific during SI, CTRL offspring = 30, PS offspring = 29, PS offspring CF to CTRL mom = 39, CTRL offspring CF to CGS mom = 39. Freezing behavior measured during (**B**) training, (**C**) context and (**D**) cued phase of FC, CTRL offspring = 30, PS offspring = 29, PS offspring CF to CTRL mom = 39, CTRL offspring CF to CGS mom = 39. Data presented as mean $\pm$ SEM. ****$p < 0.0001$ mixed linear ANOVA with prenatal stress x cross-fostering x sex model and litter as a randomized block factor.

### FIGURE 5 – FIGURE SUPPLEMENT 1

A

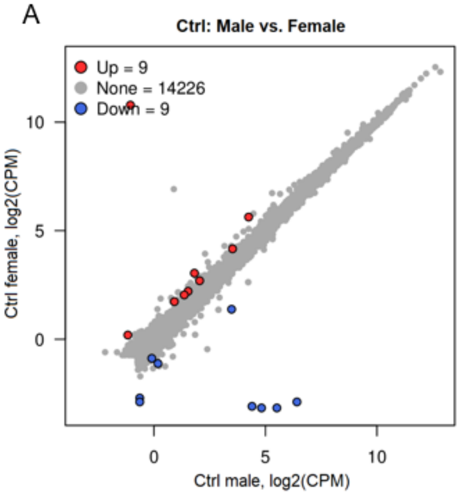


B

| Gene symbol | Chromosome | Log2FC | FDR |
|---|---|---|---|
| **Female-specific** | | | |
| Xist | X | 12.14 | 0.00E+00 |
| Prss16 | 13 | 1.99 | 1.33E-02 |
| Ntn5 | 7 | 1.39 | 2.45E-02 |
| Sec1 | 7 | 1.31 | 1.87E-04 |
| Mcmdc2 | 1 | 0.86 | 3.94E-04 |
| Pcdhb2 | 2 | 0.71 | 2.35E-02 |
| Col20a1 | 2 | 0.68 | 3.66E-02 |
| Ecm2 | 13 | 0.64 | 4.68E-03 |
| Eif2s3x | X | 0.64 | 1.19E-06 |
| **Male-specific** | | | |
| Eif2s3y | Y | -11.60 | 1.03E-268 |
| Kdm5d | Y | -11.14 | 9.20E-288 |
| Ddx3y | Y | -10.81 | 0.00E+00 |
| Uty | Y | -9.84 | 0.00E+00 |
| S100a9 | 3 | -3.19 | 1.19E-09 |
| S100a8 | 3 | -2.99 | 7.58E-09 |
| Prx | 7 | -2.28 | 5.58E-03 |
| Gm35612 | | -1.52 | 6.61E-06 |
| S100a4 | 3 | -0.89 | 3.25E-02 |

C

| GO Category ID | GO Category Name | Adjusted P-value | Hit in Query List |
|---|---|---|---|
| **Male-specific** | | | |
| GOTERM_MF:0050786 | RAGE receptor binding | 1.18E-05 | S100A4; S100A9; S100A8 |
| GOTERM_MF:0036041 | Long-chain fatty acid binding | 5.61E-03 | S100A9; S100A8 |

**Figure 5 – figure supplement 1. Analysis of DEGs in the amygdala in CTRL females vs. CTRL males.** (**A**) Scatter plot displaying 9 female-specific and 9 male-specific genes in CTRL females vs. CTRL males comparison. (**B**) Summary of DEGs. Genes were considered significant with an FDR < 0.05 and FC > 1.5. (**C**) Significantly enriched pathways after gene ontology analysis of male-specific genes using molecular function category. Genes significantly expressed in the amygdala of CTRL males when compared to CTRL females include genes encoding for calcium binding proteins associated with RAGE receptor binding process and long chain fatty acid binding process.

### FIGURE 7 – FIGURE SUPPLEMENT 1

A

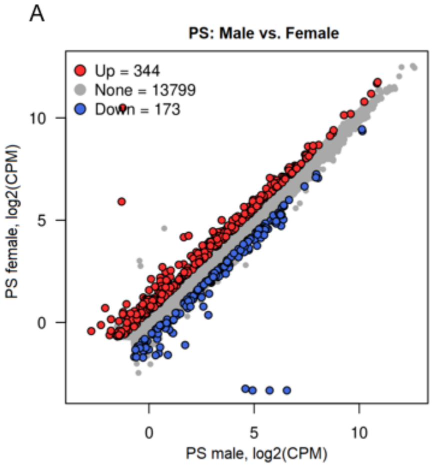


B

| GO Category ID | GO Category Name | Adjusted P-value | Hit in Query List |
|---|---|---|---|
| **PS female-specific** | | | |
| GOTERM_BP:0007268 | Synaptic transmission | 2.05E-04 | HCN4; KCNG1; CAMK2D; DOC2A; RASGRF2; DBH; SLC6A2; KCNV1; RIMS1; NPY; NPTX2; DLGAP2; CACNG3; NTSR1; GRINA3; SYT5; KCNJ4; UNC13C; KCNF1; KCNH4; KCNH5; GABRA3; SST; HAP1; CNIH3; STX1A |
| GOTERM_BP:0034762 | Regulation of transmembrane transport | 4.58E-04 | KCNG1; HCN4; KCNJ4; KCNF1; MEF2C; KCNH4; CAMK2D; KCNH5; RASGRF2; ATP2B4; TRDN; CACNG6; KCNV1; BPIFA1; JSRP1; GSG1L; HAP1; NOS1; CNIH3; CACNG3; CACNG5 |
| GOTERM_BP:1900449 | Regulation of glutamate receptor signaling | 1.35E-02 | MEF2C; RASGRF2; GSG1L; CACNG3; CNIH3; CACNG5 |
| GOTERM_CC:1902495 | Transmembrane transporter complex | 2.20E-03 | GABRQ; KCNG1; KCNJ4; KCNF1; KCNH4; CAMK2D; GABRA3; ANO2; TRDN; CACNG6; OLFM1; KCNV1; GLRA3; CNIH3; CACNG3; GRIA3; CACNG5 |

**Figure 7 – figure supplement 1. Analysis of DEGs in the amygdala in PS males vs. PS females.** (**A**) Scatter plot displaying 344 PS female-specific and 173 PS male-specific genes. Genes were considered significant with an FDR < 0.05 and FC > 1.5. (**B**) Significantly enriched pathways after gene ontology analysis of PS female-specific genes using biological process and cellular component categories.

### FIGURE 8 – FIGURE SUPPLEMENT 1

| | | RNA-seq results | | qPCR results | | | |
|---|---|---|---|---|---|---|---|
| Gene symbol | Gene name | Log2FC | FDR | FC | Log2FC | Concordant | Normalization after cross-fostering |
| **MALES** | | | | | | | |
| **Glutamatergic signaling** | | | | | | | |
| HOMER 3 | Homer Scaffold Protein 3 | 0.99 | 1.22E-02 | 3.37 | 1.75 | yes | Rescued in PS males CF to CTRL mom |
| GRM2 | Glutamate metabotropic receptor 2 | 0.75 | 1.22E-02 | 1.61 | 0.69 | yes | Rescued in PS males CF to CTRL mom |
| GSG1L | Germ cell-specific gene 1-like protein | -0.59 | 4.09E-02 | 0.57 | -0.80 | yes | |
| GRIN3A | Glutamate Ionotropic Receptor NMDA Type Subunit 3A | -0.65 | 1.21E-04 | 0.65 | -0.62 | yes | |
| CNIH3 | Cornichon Family AMPA Receptor Auxiliary Protein 3 | -0.76 | 1.47E-03 | 0.58 | -0.79 | yes | |
| CACNG5 | Calcium Voltage-Gated Channel Auxiliary Subunit Gamma 5 | -0.92 | 2.72E-04 | 0.35 | -1.51 | yes | Trend towards rescue in PS males CF to CTRL mom |
| **GABAergic signaling** | | | | | | | |
| GABRA3 | Gamma-Aminobutyric Acid Type A Receptor Subunit Alpha3 | -1.25 | 3.76E-03 | 0.61 | -0.72 | yes | |
| HAP1 | Huntingtin Associated Protein 1 | -0.85 | 1.54E-02 | 0.53 | -0.93 | yes | Trend towards rescue in PS males CF to CTRL mom |
| GABRQ | Gamma-Aminobutyric Acid Type A Receptor Subunit Theta | -0.73 | 1.58E-02 | 0.27 | -1.89 | Trend (P=0.0837) | |
| **Calcium (Ca2+) channels** | | | | | | | |
| CACNG6 | Calcium Voltage-Gated Channel Auxiliary Subunit Gamma 6 | -1.28 | 2.02E-03 | 0.16 | -2.67 | yes | |
| **Potassium (K+) channels** | | | | | | | |
| KCNA3 | Potassium Voltage-Gated Channel Subfamily A Member 3 | -0.60 | 2.82E-04 | 0.82 | -0.29 | yes | |
| KCNH4 | Potassium Voltage-Gated Channel Subfamily H Member 4 | -0.92 | 3.90E-03 | 0.51 | -0.98 | yes | |
| **FEMALES** | | | | | | | |
| DBH | Dopamine Beta Hydroxylase | 1.48 | 5.94E-03 | 4.04 | 2.01 | yes | Rescued in PS females CF to CTRL mom |
| SLC6A5 | Sodium and Chloride-dependent Glycine Transporter | 6.52 | 5.94E-03 | 3.77 | 1.92 | Trend (P=0.0899) | |

**Figure 8 supplement 1**. **Comparison of fold change determined by RNA-seq and qPCR for a subset of genes selected for validation of RNA-seq and summary of CF-induced changes.** Comparison of fold changes by RNA-seq and qPCR shows a positive correlation. Summary of gene expression changes obtained after CF obtained via qPCR has also been provided.
